# Moiety Modeling Framework for Deriving Moiety Abundances from Mass Spectrometry Measured Isotopologues

**DOI:** 10.1101/595348

**Authors:** Huan Jin, Hunter N.B. Moseley

## Abstract

**Background:** Stable isotope tracing can follow individual atoms through metabolic transformations through the detection of the incorporation of stable isotope within metabolites. This resulting data can be interpreted in terms related to metabolic flux. However, detection of a stable isotope in metabolites by mass spectrometry produces a profile of isotopologue peaks that requires deconvolution to ascertain the localization of isotope incorporation.

**Results:** To aid the interpretation of the mass spectroscopy isotopologue profile, we have developed a moiety modeling framework for deconvoluting metabolite isotopologue profiles involving single and multiple isotope tracers. This moiety modeling framework provides facilities for moiety model representation, moiety model optimization, and moiety model selection. The moiety_modeling package was developed from the idea of metabolite decomposition into moiety units based on metabolic transformations, i.e. a moiety model. A SAGA-optimize package, solving a boundary-value inverse problem through a combined simulated annealing and genetic algorithm, was developed for model optimization. Additional optimization methods from the Python scipy library are utilized as well. Several forms of the Akaike information criterion and Bayesian information criterion are provided for selecting between moiety models. Moiety models and associated isotopologue data are defined in the JSON format.

By testing the moiety modeling framework on the timecourses of ^13^C isotopologue data for UDP-N-acetyl-D-glucosamine (UDP-GlcNAc) in human prostate cancer LnCaP-LN3 cells, we were able to confirm its robust performance in isotopologue deconvolution and moiety model selection.

**Conclusions:** SAGA-optimize is a useful Python package for solving boundary-value inverse problems, and the moiety_modeling package is an easy-to-use tool for mass spectroscopy isotopologue profile deconvolution involving single and multiple isotope tracers. Both packages are freely freely available on GitHub and via the Python Package Index.

## Introduction

Recent work indicates that many human diseases involve metabolic reprogramming that disturbs normal physiology and causes serious tissue dysfunction^1^. Advances in analytical technologies, especially mass spectroscopy (MS) and nuclear magnetic resonance (NMR), have made metabolic analysis of human diseases a reality^2^. Stable isotope tracing is a powerful technique that enables the tracing of individual atoms through metabolic pathways. Stable isotope-resolved metabolomics (SIRM) uses advanced MS and NMR instrumentation to analyze the fate of stable isotopes traced from enriched precursors to metabolites, providing richer metabolomics datasets for metabolic flux analyses. NMR can measure isotopomer-specific metabolite data, but is typically limited by sensitivity. Often a single piece of NMR data only provides information on the presence of stable isotopes in just a part of a metabolite, which represents a partial isotopomer. In some cases, multiple partial isotopomer information can be interpreted in terms of a full isotopomer. MS can measure isotopologue-specific data; however, an isotopologue represents a set of mass-equivalent isotopomers. Comprehensive metabolic analysis often relies on MS metabolic datasets or a combination of MS and NMR metabolic datasets. Even though large amounts of metabolomics datasets have been generated recently, it is still a big challenge to acquire meaningful biological interpretation from MS raw data, especially for complex metabolites composed of multiple subunits or moieties.

To better interpret the complex isotopologue profile of large composite metabolites, both quantitative analysis as well as complex modeling are required. Several methods have been developed for quantitative flux analysis of specified pathways based on the stable isotope incorporated data, like the elementary metabolite units (EMU) framework^3^. These methods rely heavily on well-curated metabolic networks to accomplish the metabolic flux analysis. However, models of cellular metabolism, even for human, are far from complete.

To deconvolute the relative fluxes of complex metabolites, first a plausible model should be built based on a relevant metabolic network, which is often incomplete. When multiple models are plausible, development of a robust model selection method is essential for successful isotopologue deconvolution, especially for non-model organisms. This basic approach to isotopologue deconvolution was demonstrated in a prototype Perl program called GAIMS for the metabolite UDP-GlcNAc using a MS isotopologue profile derived from a prostate cancer cell line^4,5^. This demonstration derived relative fluxes for several converging biosynthetic pathways of UDP-GlcNAc under non-steady-state conditions. This demonstration also inspired the development of MAIMS, a software tool for metabolic tracer analysis^6^, which further validates the robustness of the moiety model deconvolution method. However, the MAIMS software handles only ^13^C single isotope tracer data and does not address model selection, which is crucial for addressing incomplete knowledge of cellular metabolic networks.

In addition, the simultaneous use of multiple stable isotopes in SIRM experiments can provide much more data than a single tracer. However, incorporation of multiple stable isotopes also complicates the analysis of metabolite isotopologue profiles, which limits most of the current isotope tracer experiments to a single tracer. The lack of data analysis tools greatly impedes the application of the multiple-labeled SIRM experiments. Therefore, we have developed a new moiety modeling framework for deconvoluting MS isotopologue profiles for both single and multiple-labeled SIRM MS datasets. This moiety modeling framework not only solves the non-linear deconvolution problem, but also selects the optimal model describing the relative fluxes of specific metabolite from a set of plausible models.

## Implementation

### Overview of the moiety modeling framework

The workflow of the moiety modeling framework is composed of three steps, model representation, model optimization, and model selection. For the model representation step, the moiety_modeling package creates an internal representation of a moiety model from a given JSONized moiety model description (see Additional file 1). In this representation illustrated by a unified modeling language (UML) class diagram in Figure 1, the package first dissembles a complex metabolite into a list of moieties, i.e. metabolic subunits. Each moiety may contain different number of labeling isotopes, representing the flow of isotope from the labeling source to the moiety. A moiety with a specific number of labeled isotopes is represented as an isotope enrichment state of the moiety (i.e. moiety state). Furthermore, mathematical relationships may exist between moiety states, even from different moieties and/or molecules. Molecules, their moieties, the possible moiety states, and relationships between moiety states work together to represent a particular moiety model, and the proportion for each possible moiety state is an optimizable parameter of the model. The next major step, moiety model optimization, involves deriving an optimal set of model parameters, i.e. moiety state fractional abundances that derives relative isotopologue abundances that best match experimental isotopologue profiles. The moiety_modeling package implements several optimization methods, including a combined simulated annealing and genetic algorithm (SAGA) based on the ‘Genetic Algorithm for Isotopologues in Metabolic Systems’ (GAIMS) Perl implementation^4,5^, a truncated Newton algorithm (TNC)^7^, a SLSQP algorithm using Sequential Least Squares Programming^8^, and a L-BFGS-B algorithm^9^. For the latter three algorithms ‘TNC’, ‘SLSQP’, and ‘L-BFGS-B’, the moiety_modeling package uses the implementation from the scipy.optimize Python module. In addition, we have the option to optimize the datasets together or separately. The final major step, model selection, tries to find the model that best fits the experimental isotopologue profiles from a set of provided moiety models. Several forms of the Akaike information criterion (AIC)^10^ and Bayesian information criterion (BIC)^11^ are used as the estimator of the relative quality of moiety models for the set of isotopologue data.

**Figure 1.**
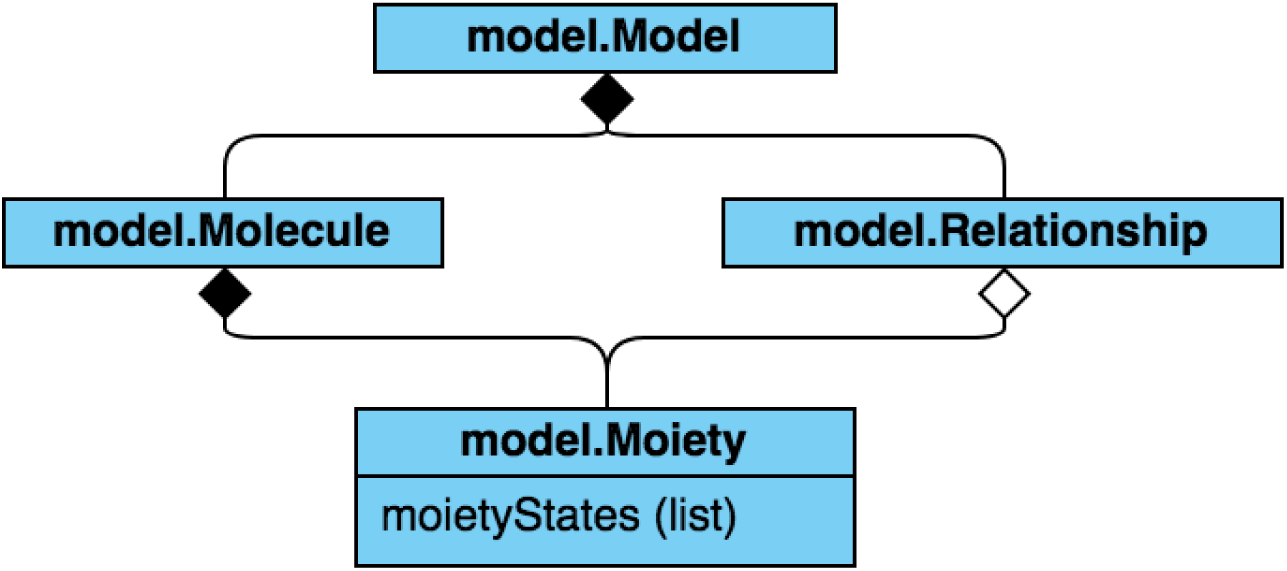
A unified modeling language (UML) class diagram of a Moiety Model.

### The moiety_modeling Python package implementation

As shown in Figure 2, the moiety_modeling Python package consists of several modules: ‘model.py’, ‘modeling.py’, ‘analysis.py’, and ‘cli.py’. The ‘model.py’ module contains class definitions for the basic elements in the moiety model. It is composed of ‘Moiety’, ‘Relationship’, ‘Molecule’ and ‘Model’ classes. The ‘Moiety’ object represents a specific moiety, the labeling isotopes present in the moiety, and their corresponding states within the moiety. The ‘Relationship’ class describes the mathematical dependencies between moiety states. A ‘Molecule’ object represents an individual metabolite made up of a list of ‘Moiety’ objects. The ‘Model’ class simulates the flow of isotope from labeling sources into each moiety of specific metabolites, which is initialized by lists of ‘Moiety’ objects, ‘Molecule’ objects, and ‘Relationship’ objects. A moiety model is generated and stored in a JSONized representation using the jsonpickle Python package^12^. This JSONized representation (see Additional file 2), stored in a file, is then used as the input file for later model optimizations. The ‘modeling.py’ module is responsible for model optimization. It is composed of the ‘Dataset’ class, several model optimization classes, and the ‘OptimizationManager’ class. The ‘Dataset’ class organizes a single MS isotopologue profile dataset into a dictionary-based data structure. ‘Dataset’ objects are stored in a JSONized representation (see Additional file 3) and used as the input for later model optimizations. There are four specific model optimization classes in the ‘modeling’ module that utilize different optimization methods and approaches for combining datasets. The ‘SAGAoptimization’ and ‘SAGAseparateOptimization’ classes use the SAGA-optimize Python package described in the next section for either combined optimization of model parameters across all datasets or separate optimizations of model parameters for each dataset. ‘ScipyOptimization’ and ‘ScipySeparateOptimization’ classes make use of optimization methods (‘TNC’, ‘SLSQP’, and ‘L-BFGS-B’) in the scipy.optimize module to conduct optimizations in either a combined or separate manner. The ‘OptimizationManager’ class is responsible for the management of the optimization process based on the input optimization parameters. The results for a model optimization are stored in a JSONized representation (see Additional file 4) for further analysis. A text file is used to store the filepaths to all of the optimized models with certain optimization parameters. The filepath file is then used as the input for the ‘analysis.py’ module. The ‘analysis.py’ module has five classes: ‘ResultsAnalysis’, ‘ModelRank’, ‘ComparisonTable’, ‘PlotMoietyDistribution’ and ‘PlotIsotopologueIntensity’. The ‘ResultsAnalysis’ class is responsible for generating standard statistics from the optimization results for each model. The mean, standard deviation, minimum, and maximum value of each model parameter are calculated. The calculated isotopologue intensities and their statistics based on the sets of optimized parameters are also generated. Furthermore, several quality estimators of each model, including different forms of the ‘AIC’ (Table 1), are computed for model selection. The AIC tends to select the model that has too many parameters when the sample size is small, leading to overfitting. The sample size corrected AIC (AICc) was developed to address this overfitting problem^13^. The Bayesian information criterion (BIC) is another commonly used criterion for model selection^14^. The ‘ResultsAnalysis’ objects with results for each model are stored in a JSONize representation (see Additional file 5) for further analysis, along with a text report for readability. Also, an analysis filepath file containing the filepaths to the analysis JSON files of all models with the same optimization parameters is created. Next, the ‘ModelRank’ class object uses this analysis filepath file to compare and select the model that best reflects the observed isotopologue profile. The ‘ComparisonTable’ class compares the model selection results with different optimization parameters. The ‘PlotMoietyDistribution’ class and ‘PlotIsotopologueIntensity’ class are responsible for visualization of the optimization results. The ‘cli.py’ module provides the command-line interface to perform model optimization, model optimization analysis, and model selection, which is implemented with the ‘docopt’ Python library^15^.

**Table 1.** Different forms of a model selection estimator.

| **Selection Criterion** | **Equation** |
| --- | --- |
| Akaike Information Criterion (AIC) | $2k + n\ln(RSS/n)$ |
| Sample size corrected AIC (AICc) | $AIC + (2k^2 + 2k)/(n - k - 1)$ |
| Bayesian Information Criterion (BIC) | $n\ln(RSS/n) + k\ln(n)$ |

k is the number of parameters.
n is the number of data points.
RSS is the residual sum of squares: $RSS = \sum_{i=1}^n (I_{obs} - I_{calc})^2$ .

**Figure 2.**
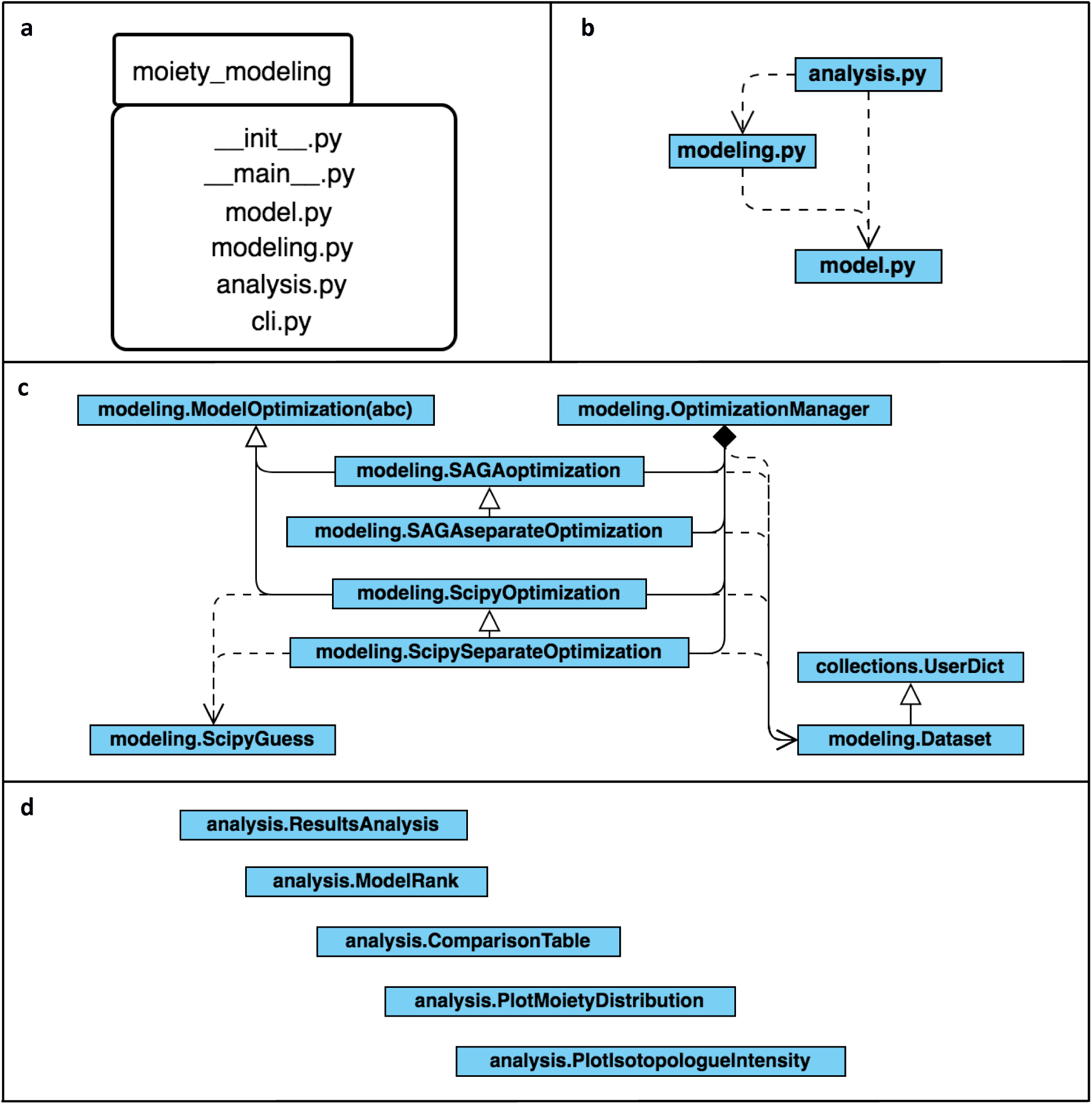
Organization of the moiety_modeling package represented with UML diagrams: **a** UML package diagram of the moiety_modeling Python library; **b** Subpackage dependencies diagram; **c** UML class diagram of the ‘modeling.py’ module with dependency relationships; **d** UML class diagram of the ‘analysis.py’ module, which contains a set of classes with no relationships.

### SAGA-optimize Python package implementation

The SAGA-optimize Python package is a novel type of combined simulated annealing and genetic algorithm^4^ used to find the optimal solutions to a set of parameters based on a given energy function calculated using the set of parameters. In this context, the energy function represents a comparison of calculated and experimentally observed isotopologue relative intensities and the calculated intensities are based on the moiety model parameters being optimized. As shown in Figure 3, it is composed of ‘ElementDescription’, ‘Guess’, ‘Population’ and ‘SAGA’ classes. An ‘ElementDesciption’ object describes an individual parameter of the moiety model. The ‘ElementDescription’ object is bound by a range and several mutation methods are available to change the value of the ‘ElementDescription’ object. A ‘Guess’ object contains lists of all the parameters (‘ElementDescription’ objects) and their corresponding values for a particular moiety model. In addition, it also stores the energy calculated based on this set of parameters. A ‘Population’ object contains information of a list of ‘ElementDescription’ objects, a list of ‘Guess’ objects, the range of each ‘ElementDescription’ among all the ‘Guess’ objects, the highest and lowest energy for the list of ‘Guess’ objects, and the best ‘Guess’ object. The ‘ElementDescription’, ‘Guess’ and ‘Population’ classes are the building blocks of the ‘SAGA’ class, which is the main class that provides the interface for optimization. Furthermore, several distinct crossover functions are available for creating new Guess objects from the cross-over of two other Guess objects.

**Figure 3.**
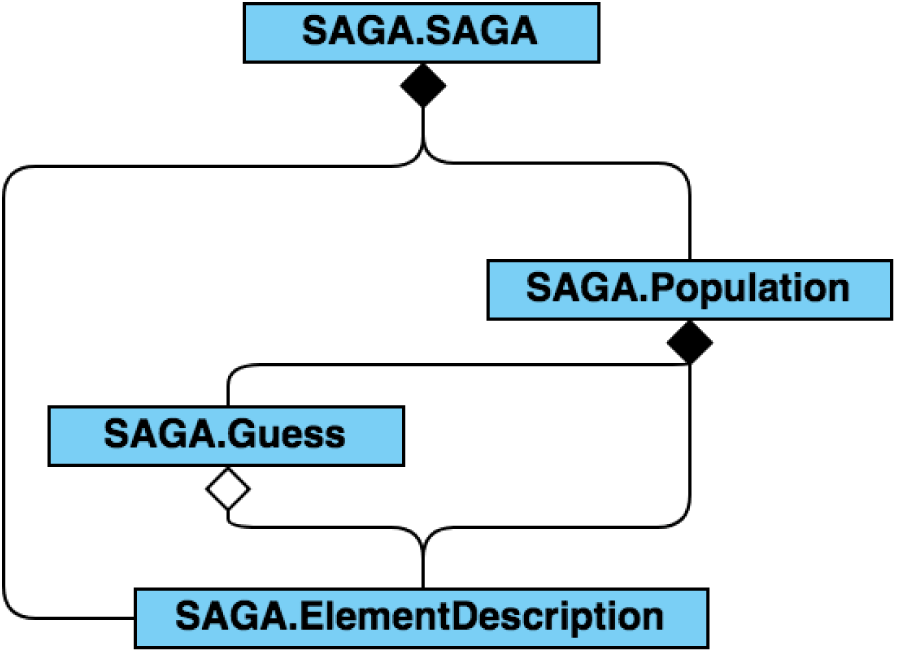
‘SAGA’ package represented with a UML class diagram with dependencies.

## Results and discussion

### The package interface

The moiety_modeling package can be used in two main ways: (i) as a library within Python scripts for accessing and manipulating moiety models and isotopologue datasets stored in JSON files, or (ii) as a command-line tool to perform model optimization, model analysis, and model selection.

To use the moiety_modeling package as a library within Python scripts, it should be imported with a Python program or an interactive interpreter interface. Next, ‘Moiety’, ‘Relationship’ and ‘Molecule’ objects can be created to construct a moiety model. ‘Dataset’ objects are also built with the moiety_modeling package. Table 2 summarizes common patterns for using moiety_modeling package as a library in construction of a moiety model and related datasets.

**Table 2.** Common creation patterns for the moiety_modeling library.

| Entity | Example |
| --- | --- |
| Moiety | glucose = moiety_modeling.Moiety('glucose', {'13C': 6}, {'13C': [1, 3, 5]}, 'g')<br>acetyl = moiety_modeling.Moiety('acetyl', {'13C': 2}, {'13C': [0, 1, 2]}, 'a')<br>uracil = moiety_modeling.Moiety('uracil', {'13C': 4}, {'13C': [1, 2, 4]}, 'u')<br>ribose = moiety_modeling.Moiety('ribose', {'13C': 5}, {'13C': [0, 3, 5]}, 'r') |
| Relationship | relationship = moiety_modeling.Relationship(glucose, '13C0', acetyl, '13C2', '*', 2) |
| Molecule | UDP-GlcNAc = moiety_modeling.Molecule('UDP-GlcNAc', [glucose, uracil, acetyl, ribose]) |
| Model | model1 = moiety_modeling.Model('model1', [glucose, uracil, acetyl, ribose], [UDP_GlcNAc], [relationship]) |
| Dataset | dataset = moiety_modeling.Dataset('12h', 'UDP_GlcNAc': [{'labelingIsotopes': '13C0', 'height': 0.0175, 'heightSE': 0 }, {'labelingIsotopes': '13C1', 'height': 0.0075, 'heightSE': 0 }, ...] ) |

The moiety_modeling package also provides a simple command-line interface to perform model optimization, selection, and visualization. Additional file 6 shows version 1.0 of the command-line interface, and Table 3 summarizes common pattern for using moiety_modeling as a command-line tool. The common patterns for using SAGA-optimize as a library are shown in Additional file 7.

**Table 3.** Common patters for using the moiety_modeling as a command-line tool.

| Command | Description | Example |
| --- | --- | --- |
| modeling | Perform model optimization | % python3 -m moiety_modeling modeling --models=models.json --datasets=dataset.json --optimizations=optimization_settings.json |
| analyze | Analyze the optimization results | % python3 -m moiety_modeling analyze optimizations --a optimizationPaths.txt |
| plot | Plot the distribution of calculated moiety modeling parameters. | % python3 -m moiety_modeling plot moiety analysisResults.json |

### Advantage of JSONized representation for MS isotopologue data and analysis results

JavaScript object notation (JSON)^16^ is an open-standard file format using human-readable text to collect data in pair-value and array structures, widely used by different programming language. Complex Python objects, like ‘Moiety’ and ‘Molecule’ objects mentioned above, can be serialized to JSON format with the jsonpickle Python library. The moiety model and dataset constructed with moiety_modeling package as well as optimization parameters are the input files for the moiety modeling, all of which are saved in JSON format using jsonpickle (see Additional file 2, 3, and 8). The use of JSON format makes the moiety modeling framework easily accessible to other programming languages and naturally extendible. In addition, the optimization and analysis results are also stored in a JSON file (see Additional file 4 and 5).

### Dataset and model

We used the timecourse of ^13^C isotopologue data for UDP-GlcNAc generated from [U-^13^C]-glucose in human prostate cancer LnCaP-LN3 cells to evaluate the robustness of the moiety modeling framework. 40 hypothetical moiety models of the isotopic flow into UDP-GlcNAc were crafted manually. Also, an expert-derived moiety model of UDP-GlcNAc was created based on known biochemical pathways and corroborated by NMR data. We tested whether the expert-derived moiety model could be selected from all the other models.

### Data validation

The incorporation of ^13^C from [U-^13^C]-glucose into UDP-GlcNAc leads to a total of 17 isotopologues. We applied the moiety modeling framework to the observed UDP-GlcNAc isotopologue data with each built model to test whether the expert-derived moiety model could be selected above the others. The results are list in the Table 4. From the results, we can see that the expert-derived moiety model can be selected successfully among all the moiety models, which demonstrates the robustness of the moiety modeling framework.

**Table 4.**
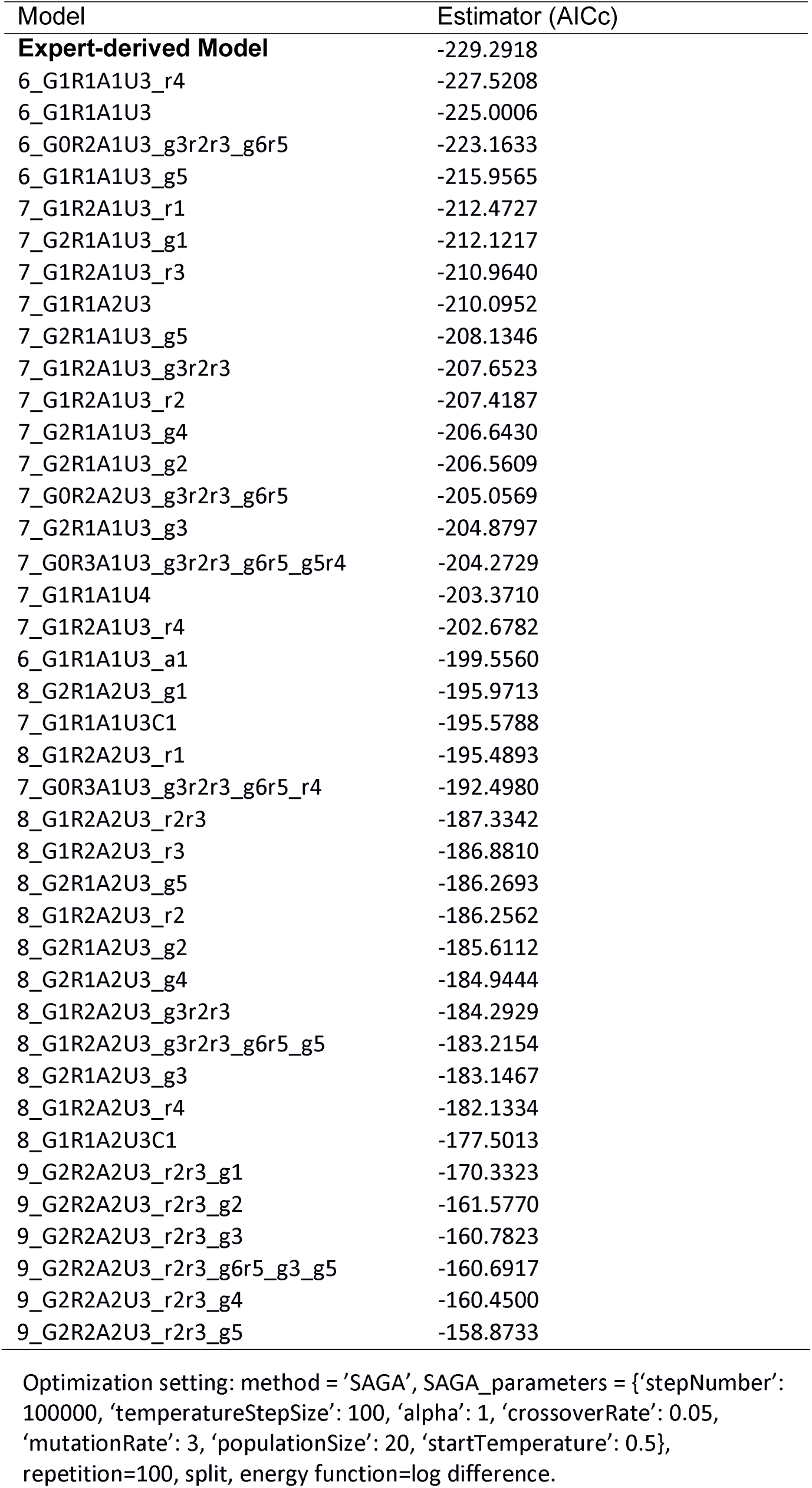
Model selection results of UDP-GlcNAc isotopologue data

| Model | Estimator (AICc) |
| --- | --- |
| **Expert-derived Model** | -229.2918 |
| 6_G1R1A1U3_r4 | -227.5208 |
| 6_G1R1A1U3 | -225.0006 |
| 6_G0R2A1U3_g3r2r3_g6r5 | -223.1633 |
| 6_G1R1A1U3_g5 | -215.9565 |
| 7_G1R2A1U3_r1 | -212.4727 |
| 7_G2R1A1U3_g1 | -212.1217 |
| 7_G1R2A1U3_r3 | -210.9640 |
| 7_G1R1A2U3 | -210.0952 |
| 7_G2R1A1U3_g5 | -208.1346 |
| 7_G1R2A1U3_g3r2r3 | -207.6523 |
| 7_G1R2A1U3_r2 | -207.4187 |
| 7_G2R1A1U3_g4 | -206.6430 |
| 7_G2R1A1U3_g2 | -206.5609 |
| 7_G0R2A2U3_g3r2r3_g6r5 | -205.0569 |
| 7_G2R1A1U3_g3 | -204.8797 |
| 7_G0R3A1U3_g3r2r3_g6r5_g5r4 | -204.2729 |
| 7_G1R1A1U4 | -203.3710 |
| 7_G1R2A1U3_r4 | -202.6782 |
| 6_G1R1A1U3_a1 | -199.5560 |
| 8_G2R1A2U3_g1 | -195.9713 |
| 7_G1R1A1U3C1 | -195.5788 |
| 8_G1R2A2U3_r1 | -195.4893 |
| 7_G0R3A1U3_g3r2r3_g6r5_r4 | -192.4980 |
| 8_G1R2A2U3_r2r3 | -187.3342 |
| 8_G1R2A2U3_r3 | -186.8810 |
| 8_G2R1A2U3_g5 | -186.2693 |
| 8_G1R2A2U3_r2 | -186.2562 |
| 8_G2R1A2U3_g2 | -185.6112 |
| 8_G2R1A2U3_g4 | -184.9444 |
| 8_G1R2A2U3_g3r2r3 | -184.2929 |
| 8_G1R2A2U3_g3r2r3_g6r5_g5 | -183.2154 |
| 8_G2R1A2U3_g3 | -183.1467 |
| 8_G1R2A2U3_r4 | -182.1334 |
| 8_G1R1A2U3C1 | -177.5013 |
| 9_G2R2A2U3_r2r3_g1 | -170.3323 |
| 9_G2R2A2U3_r2r3_g2 | -161.5770 |
| 9_G2R2A2U3_r2r3_g3 | -160.7823 |
| 9_G2R2A2U3_r2r3_g6r5_g3_g5 | -160.6917 |
| 9_G2R2A2U3_r2r3_g4 | -160.4500 |
| 9_G2R2A2U3_r2r3_g5 | -158.8733 |

Optimization setting: method = 'SAGA', SAGA\_parameters = {'stepNumber': 100000, 'temperatureStepSize': 100, 'alpha': 1, 'crossoverRate': 0.05, 'mutationRate': 3, 'populationSize': 20, 'startTemperature': 0.5}, repetition=100, split, energy function=log difference.

## Conclusions

Here, we present a moiety modeling framework for the deconvolution of metabolite isotopologue profiles using moiety models along with the analysis and selection of the best moiety model(s) based on the experimental data. This framework can analyze datasets involving single and multiple isotope tracers. With a ^13^C-labeled UDP-GlcNAc isotopologue dataset, we demonstrate the robust performance of the moiety modeling framework for model selection. The selection of correct moiety models is required for generating deconvolution results that can be accurately interpreted in terms of relative metabolic flux. Furthermore, the JSON formats of moiety model, isotopologue data, and optimization results facilitate the inclusion of these tools in data analysis pipelines.

## Availability and requirements

**Project name:** moiety_modeling

**Pipeline Installation manual:** https://moiety-modeling.readthedocs.io/en/latest/

**Operating system:** Linux

**Programming language:** Python 3.5+

**Other requirements:** jsonpickle, matplotlib, docopt, scipy, numpy

**License:** BSD

## Additional files

**Additional file 1:** JSONized description of moiety model components. (DOCX)

**Additional file 2:** Moiety model description. (JSON)

**Additional file 3:** Dataset for moiety modeling. (JSON)

**Additional file 4:** Optimization results. (JSON)

**Additional file 5:** Analysis results. (JSON)

**Additional file 6:** The moiety_modeling package command line interface (DOCX)

**Additional file 7:** Common patterns for using ‘SAGA’ module as a library (DOCX)

**Additional file 8:** Optimization parameters. (JSON)

### Abbreviations

MS: Mass spectroscopy
NMR: Nuclear magnetic resonance
SIRM: Stable isotope-resolved metabolomics
UDP-GlcNAc: UDP-N-acetyl-D-glucosamine
SAGA: Simulated annealing and genetic algorithm
GAIMS: Genetic Algorithm for Isotopologues in Metabolic Systems
AIC: Akaike information criterion
BIC: Bayesian information criterion
UML: Unified modeling language
JSON: JavaScript Object Notation

## Declarations

## Availability of data and materials

The moiety_modeling and SAGA-optimize packages are available on: GitHub - https://github.com/MoseleyBioinformaticsLab/moiety_modeling, https://github.com/MoseleyBioinformaticsLab/SAGA_optimize. PyPI - https://pypi.org/project/moiety-modeling/, https://pypi.org/project/SAGA-optimize/.

Project documentation is available online at ReadTheDocs: https://moiety-modeling.readthedocs.io/en/latest/, https://saga-optimize.readthedocs.io/en/latest/.

All the results analyzed in this manuscript are available on figshare: https://figshare.com/articles/moiety_modeling_framework/7886135.

## Author’s contribution

HJ and HNBM worked together on the design of the Python libraries, their API, and their CLI (moiety_modeling only). HJ implemented the libraries. HNBM helped troubleshoot implementation issues and redesign. HJ and HNBM wrote the manuscript. All authors have read and approved the manuscript.

## Consent for publication

Not applicable.

## Ethics approval and consent to participate

Not applicable

