## Supplementary material for "Moiety Modeling Framework for Deriving Moiety Abundances from Mass Spectrometry Measured Isotopologues": Additional_file_1.docx

**Moiety**

{

“name”: “ribose”,

“nickname”: “r”,

“ranking”: 1,

“maxIsotopeNum”: {“13C”: 5},

“isotopeStates”: {“13C”: [0, 2, 3, 5] },

“states”: [ “13C0”, “13C2”, “13C3”, “13C5” ] ,

“py/object”: “moiety_modeling.model.Moiety”

}

**Molecule**

{

“name”: “UDP_GlcNAC”,

“moieties”: [{“py/id”: 13}, …],

“standardStates”: {“13C0”: [ [ “ribose[13C0]”, “glucose[13C0]”, “acetyl[13C0]”,

“uracil[13C0]”]], “13C10”: [ [“ribose[13C0]”, “glucose[13C6]”, “acetyl[13C2]”,

“uracil[13C2]”], [“ribose[13C5]”, “glucose[13C0]”, “acetyl[13C2]”,

“uracil[13C3]”], …], …},

“allStates”: [ “13C0”, “13C1”, “13C2”, “13C3”, “13C4”, “13C5”, “13C6”, “13C7”, “13C8”,

“13C9”, “13C10”, “13C11”, “13C12”, “13C13”, “13C14”, “13C15”, “13C16”, “13C17”],

“py/object”: “moiety_modeling.model.Molecule”

}

**Relationship**

{

“moiety”: {“py/id”: 13},

“moietyState”: “13C0”,

“varName”: “glucose[13C0]”,

“equivalentMoiety”: {“py/id”: 10},

“equivalentMoietyState”: “13C0”,

“equivalentVarName”: “ribose[13C0]”,

“multiplier”: 1,

“py/object”: “moiety_modeling.model.Relationship”

}

JSONized moiety model description.
