## Supplementary material for "Moiety Modeling Framework for Deriving Moiety Abundances from Mass Spectrometry Measured Isotopologues": Additional_file_6.docx

The moiety_modeling command-line interface.

Usage:

moiety_modeling –h | --help

moiety_modeling --version

moiety_modeling modeling [--combinedData=<combined_jsonfile>] [--models=<models_jsonfile>] [--datasets=<datasets_jsonfile>] [--optimizations=<optimizations_jsonfile>] [--working=<working_dir>] [--repetition=<optim_count>] [--split] [--force] [--multiprocess] [--energyFunction=<function>] [--printOptimizationScripts]

moiety_modeling analyze optimization --a <optimzationPaths_txtfile> [--working=<working_dir]

moiety_modeling analyze optimization --s <optimzationResults_jsonfile> [--working=<working_dir]

moiety_modeling analyze rank <analysisPaths_txtfile> [--working=<working_dir>] [--rankCriteria=<rankCriteria>]

moiety_modeling analyze table <rankPaths_txtfile> [--working=<working_dir>]

moiety_modeling plot moiety <analysisResults_jsonfile> [--working=<working_dir>]

moiety_modeling plot isotopologue <analysisResults_jsonfile> [--working=<working_dir>]

Options:

-h, --help	Show this screen.

--version	Show version.

-combinedData JSON description file of the combined data (eg: models, datasets, optimization settings)

--models=<models_jsonfile>	JSON description file of the moiety models.

--datasets=<datasets_jsonfile>	JSON description file of the datasets.

--optimizaitons=<optimizaitons_jsonfile> JSON description file of the optimization setting.

--working=<working_dir>	Alternative path to save the results.

--repetition=<optim_count> The number of optimization repetitions to perform [default: 100].

--split	To split the datasets or not.

--force	To force optimization process if error occurs.

--mulitprocess	To perform with multiprocessing or not.

--printOptimizationScripts	To print the optimization script or not.

--a	To analyze a bunch of optimization results together with the path file.

--s	To analyze a single moiety model optimization results.

--energyFunction=<function> The energy function for optimization [default: logDifference].

--optimzationSetting=<optimizationSetting> The optimization setting of the moiety modeling optimization.

--rankCriteria=<rankCriteria>	The criteria for model ranking [default: AIC]

The ‘moiety_modeling’ package command line interface.
