## Supplementary material for "Moiety Modeling Framework for Deriving Moiety Abundances from Mass Spectrometry Measured Isotopologues": Additional_file_7.docx

| **Table S1.** Common patterns for using ‘SAGA’ module as a library. | |
| --- | --- |
| **Usage** | **Example** |
| SAGA | saga = SAGA.SAGA(stepNumber=100, temperature=10, startTemperature=0.5, alpha=1, energyfunction=targertedEnergyFunction) |
|  | saga.addElmentDescriptions(SAGA.ElementDecription(low=0, high=1)) |
| Population | population = saga.optimize() |
| Guess | bestGuess = population.bestGuess |
